# Quantification of STLV-1 *Tax* in various tissues of non-human primates by different PCR methods

**DOI:** 10.64898/2026.07.27.739517

**Authors:** Alexandra Birzer, Melissa Kiessling, Alina Russ, Slaveia Garbit, Valérie Moulin, Marie Isnardon, Lucie Faccin, Alexia Cermolacce, Sandrine Alais, Hélène Dutartre, Chloé Journo, Andrea K Thoma-Kress

## Abstract

**Background:** Human T-lymphotropic virus type 1 (HTLV-1) is an oncogenic retrovirus which is transmitted via cell-containing blood fluids or from mother to child via breastfeeding, leading to severe diseases such as adult T-cell leukemia/lymphoma (ATL) and neuroinflammation. Most studies focus on virus detection in peripheral blood due to limited access of tissue material, especially in infants. Thus, our understanding of viral distribution in organs, in particular along the oral route of transmission, is still a critical gap in HTLV-1 research.

**Methodology/Principal Findings:** Here, we present an analysis of tissues from a non-human primate (NHP) colony (olive baboon: *Papio anubis*) naturally infected with the closely related counterpart of HTLV-1, simian T-lymphotropic virus type 1 (STLV-1). Various organs and tissues of the oropharyngeal and gastrointestinal tract including tonsils, stomach, small intestine and colon were analyzed for the presence or absence of STLV-1. Beside TaqMan qPCR measuring relative copy numbers, we established a highly sensitive and precise droplet digital (dd) PCR protocol to measure absolute copy numbers of viral *Tax* DNA. *Tax* DNA was detectable in the tonsils in two NHPs, but to a greater extent in stomach in four NHPs. We also found *Tax* in parts of the small intestine, *i.e.* duodenum and Peyer’s patches, in a NHP with high blood proviral load.

**Conclusion/Significance:** These data provide a quantitative analysis of STLV-1 *Tax* in the gastrointestinal tract and are, to our knowledge, the first indication of STLV-1 detection in stomach tissue of naturally STLV-1-infected asymptomatic NHPs. Although it is unclear how infection occurred - from mother-to-child, sexual or via animal bites - our study suggests that these parts of the gastrointestinal tract might either serve as site of virus transmission or as viral reservoir.

*Author summary:* Human T-lymphotropic virus type 1 (HTLV-1) is a human oncogenic retrovirus being transmitted via cell-containing body fluids such as breast milk, blood, or semen. The estimated number of infected people is around 10 to 20 million. To study the viral distribution and persistence of HTLV-1 in different organs in the oropharyngeal and gastrointestinal tract, a suitable *in vivo* model is essential. In this study, tissues of non-human primates (NHPs, baboon: *Papio anubis*) naturally infected with the closely related simian counterpart of HTLV-1, simian T-lymphotropic virus type 1 (STLV-1), were analyzed for the presence or absence of the viral gene *Tax*. We identified the presence of STLV-1 *Tax* in the stomach, duodenum and Peyer’s patches by using two different detection methods: TaqMan-based qPCR and the more sensitive droplet digital PCR (ddPCR). *Tax* could be detected in tonsils in two NHPs only, but in stomach in four NHPs. Together, this is the first time that STLV-1 *Tax* was detected in stomach tissue of STLV-1-infected asymptomatic NHPs, highlighting the importance of investigating viral persistence and/or viral reservoirs of primate T-lymphotropic viruses independently of the entry route.

## Introduction

Few viruses are transmitted from mother-to-child including human immunodeficiency virus type 1 (HIV-1), human cytomegalovirus (HCMV) and human T-lymphotropic virus type 1 (HTLV-1) (5). HTLV-1 is transmitted from cell-to-cell and predominantly integrates and persists in human CD4^+^ T cells. Next to transmission from mother-to-child, HTLV-1 also transmits via sexual contacts or exchange of cell-containing body fluids such as blood products (6). For HTLV-1, not only the entry site of viral transmission along the oral route but also viral dissemination in the oropharyngeal and gastrointestinal tract remains incompletely understood. HIV-1 studies already have proven the persistence of the virus in the gastrointestinal tract, serving as viral reservoir (7, 8). HTLV-1 represents a worldwide distributed virus which is endemic in Japan, South America, the Caribbean, sub-Saharan Africa and Central Australia (9). Within the estimated 10 to 20 million people infected with HTLV-1, around 5 % develop adult T-cell leukemia/lymphoma (ATL), a neoplasia with poor prognosis (10–12). Remarkably, if HTLV-1 is acquired during infancy, the life-time risk to develop ATL is estimated to increase up to 25 % (13). Thus, there is an urgent need to understand viral entry sites and viral reservoirs. Due to the limited availability of tissue samples, especially from infants, most studies have focused on peripheral blood. Thus, there remain significant gaps in our understanding of HTLV-1 organ distribution along the oropharyngeal and gastrointestinal tract independent of the route of viral transmission. Studying animal models infected with related primate T-lymphotropic viruses (PTLVs) is therefore a beneficial way to shed light on viral dissemination in the oropharyngeal and gastrointestinal tract.

HTLV-1 has emerged from cross-species transfer (zoonosis) from non-human primates (NHPs) to humans (14). The simian counterpart simian T-lymphotropic virus type 1 (STLV-1) infected humans in several independent events in different regions around the world, giving rise to the different subtypes of HTLV (15). Together, HTLVs and STLVs are grouped into the PTLVs. STLV-1 naturally infects a variety of Old World monkeys, such as macaques, olive baboons, African green monkeys, orangutans, chimpanzees and apes (16–18), and is distributed predominantly in Asia and Africa (19). The close relationship between HTLV-1 and STLV-1 is exemplified by a 96 % homology of STLV-1 to HTLV-1 (cosmopolitan subtype A) regarding the amino acid sequence of the regulatory protein Tax (20). Nevertheless, several STLV-1 strains isolated from different infected animal species are divergent from other PTLV-1s as characterized by their lack of coding open reading frames for certain accessory genes (1, 21). Like HTLV-1, STLV-1 persists lifelong within the host (20, 22, 23). While HTLV-1 is associated with ATL and inflammatory conditions such as HTLV-1-associated myelopathy/tropical spastic paraparesis (HAM/TSP), the latter disease is not observed in STLV-1-infected NHPs, but 3 to 4 % of STLV-1-infected NHPs, such as baboons, develop ATL-like diseases (2, 20, 24, 25). The reason for this observation is unclear. Like HTLV-1, STLV-1 is also transmitted horizontally-sexual or via animal bites, and vertically-from mother to child-via mother-to-child transmission (MTCT) (6, 26). Given the similarities between the human and simian virus strains, STLV-1 naturally infected NHPs like the olive baboon (*Papio anubis)* represent a valuable *in vivo* model to study infection of STLV-1 and viral dissemination.

For the deltaretrovirus STLV, especially STLV-2 and STLV-3, viral DNA was detected in fecal samples of wild-living bonobos suggesting involvement of the gastrointestinal tract or the gut-associated lymphoid tissue in transmission, dissemination or reservoir formation of STLVs (27). While our understanding of viral distribution in asymptomatic virus carriers is limited, gastric involvement has been well documented for the HTLV-1 associated highly infiltrative ATL, showing ulcerative lesions containing infected tumor cells (28–32). Based on a study analyzing organs of an animal with ATL, the proviral load (PVL) was mainly detected in lymph nodes, spleen and lungs (2). This suggests that the gastrointestinal tract, harboring a plethora of different immune cells, might play a role in virus dissemination and reservoir formation. However, data for STLV-1 and asymptomatic PTLV infection remain scarce.

In this study, we aimed to measure STLV-1 infection within different tissues of the oropharyngeal and gastrointestinal tract of asymptomatic naturally STLV-1-infected olive baboons (*Papio anubis*). Using two different methods, TaqMan quantitative (qPCR) and a sensitive droplet digital (dd) PCR protocol, we analyzed *Tax* DNA levels in eight animals from a naturally STLV-1-infected *Papio anubis* colony housed in a Primate Center in the south of France, including seven asymptomatic and one ATL animal. We retrospectively analyzed a biobank of tissues from these animals, which precluded tracing the route of transmission. STLV-1 infection models represent a valuable surrogate for studying dissemination of the closely-related HTLV-1. Our study shows for the first time that STLV-1 is not only detectable in the duodenum and Peyer’s patches, but also in the stomach tissue, supporting the hypothesis that the gastrointestinal tract and its associated lymphoid tissues may play a role in transmission, dissemination or reservoir formation of STLV-1.

## Methods

### Ethics statement

The animals belong to a previously described cohort (33) of naturally STLV-1-infected baboons (*Papio anubis*) housed at the Primate Center of the Centre National de la Recherche Scientifique (UAR 846) in Rousset-sur-Arc, France, and were cared for in compliance with French regulations and guidelines of the Federation of European Laboratory Animal Science Associations (FELASA). The use of animals was approved by the Ethics Committee number 14 (APAFIS #4227– 201604130940121; APAFIS #34214-2021111215342100 v4) of the French Minister of Education and Research. The experimental procedure complied with the current French laws and the European directive 86/609/CEE. Blood was obtained after anesthesia via the intramuscular injection of ketamine (4-5 mg/kg) and medetomidine (0.04-0.05 mg/kg), and tissue samples were obtained post-mortem after intravenous injection of pentobarbital (180 mg/kg).

### Animals

In the cohort of naturally STLV-1-infected baboons, STLV-1 infection was initially determined by serologic methods (immunofluorescence with HTLV-1 infected MT-2 cells or ELISA performed in the Biomedical Primate Research Centre, The Netherlands) and confirmed for some of them by Western blotting (MP diagnostics HTLV-1/II 2.4). Eight STLV-1-infected and two non-infected female NHPs were included in the present study (Table 1). Some of these had already been included and described in previous studies (2–4, 33). The proviral load (PVL) was measured in whole blood and in peripheral blood mononuclear cells (PBMC) as described previously (2, 3). In the course of past studies, some STLV-1-infected animals had received treatment with antiretrovirals (Table 1), such as the nucleoside analogue Zidovudine (3’-azido-3’-deoxythymidine; AZT) or Viread (GILEAD Science GmbH), which contains Tenofovir disoproxil. Antiretrovirals were combined with the epigenetic regulator valproic acid (VPA) (33) and/or with alpha interferon (Table 1) (2). Among the STLV-1-infected animals, we included one animal who was diagnosed with ATL and euthanized in 2014 with a very high PVL of 1.1*10^5^ copies/10^5^ cells (NHP ATL) (2). The seven other STLV-1-infected animals remained lifelong asymptomatic with PVLs ranging from 3.7*10^1^ to 6.8*10^3^ copies/10^5^ cells as measured in 2016 in PBMCs or peripheral blood, and were euthanized for reasons independent of STLV-1 infection in 2019, 2020 or 2022 (Table 1). Co-housing groups and kinship among animals are also indicated in Table 1.

**Table 1:**
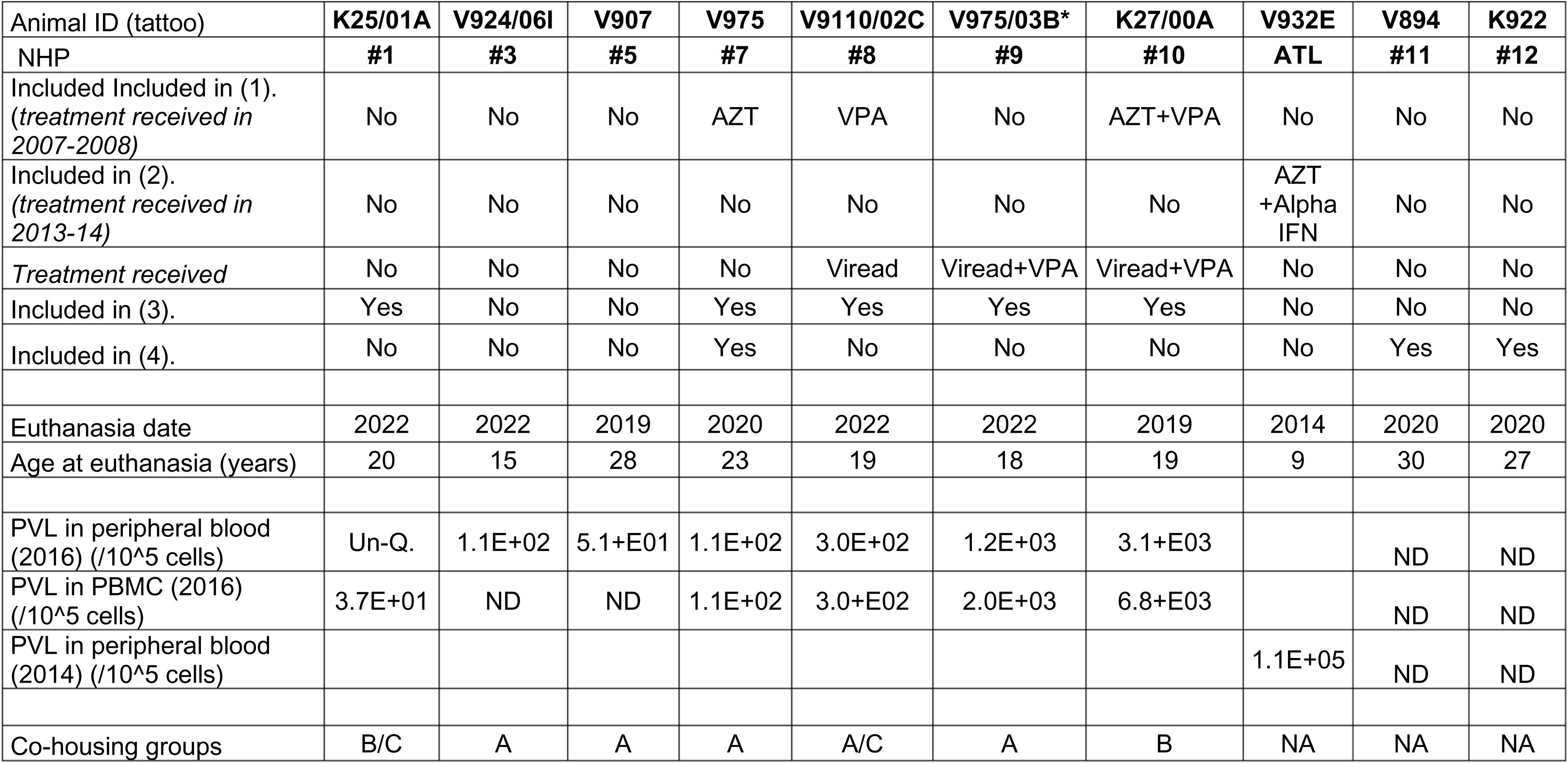
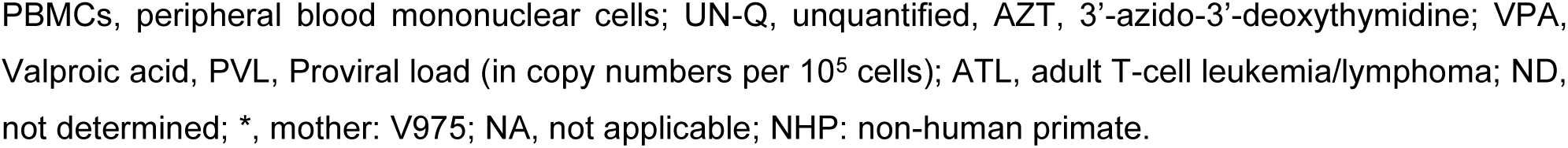
NHP cohort.

### Cell culture

HTLV-1-infected C91-PL cells were cultivated in RPMI 1640 (Gibco, Life Technologies, Darmstadt, Germany), 10 % fetal bovine serum (FBS, Anprotec, Bruckberg, Germany), 0.12 mg/ml Penicillin, 0.12 mg/ml Streptomycin and 1 % GlutaMax (all Gibco). Jurkat cells were cultivated in a 1:1 mixture of RPMI 1640 and Panserin (PAN, Biotech, Germany) supplemented with FBS, Penicillin, Streptomycin and GlutaMax like C91-PL cells. Every second to third day, cells were split in a ratio of 1:3 in media and cultivated at 37 °C and 7 % CO_2_. Authenticity of cell lines was verified by short tandem repeat analysis (DSMZ, Braunschweig, Germany).

### Tissue lysis and DNA isolation

Samples from different organs of the oropharyngeal and gastrointestinal tract, including the tonsils, stomach, intestine (and more specifically the jejunum, duodenum, Peyer’s patches, caecum, appendix and ascending colon), were snap frozen after surgical excision and stored at -80°C. Samples were cut using a scalpel in a petri dish and transferred into Precellys 2 mL Reinforced Tubes containing 1.44 mm Precellys Ceramic Beads (Bertin Instruments, Montigny-le-Bretonneux, France). After addition of 180 µL freshly-prepared lysis buffer and 25 µL proteinase K, according to the manufacturer’s instructions of the NucleoSpin Tissue Mini kit (Macherey and Nagel, Düren, Germany), samples were vortexed and incubated for 30 sec at 6,500 rpm in the Precellys 24 Tissue Homogenizer for tissue disruption. Afterwards, samples were centrifuged for 4 min at 8,609 g. Subsequent steps for DNA isolation for tissue and cells were performed as described by the manufacturer (NucleoSpin Tissue Mini kit). DNA was eluted in 50 µL elution buffer, and concentration was measured at NanoDrop1000 spectrophotometer (Thermo Fisher Scientific, Massachusetts, USA).

### TaqMan quantitative real-time PCR (qPCR)

For the TaqMan qPCR, 200 ng DNA (in 10 µL) were mixed with a master mix containing 12.5 µL of TaqMan Universal PCR Master Mix (1-fold, Life Technologies, California, USA), 0.2 µM of forward (fwd, 0.5 µL) or reverse (rev, 0.5 µL) primer, 0.25 µM TaqMan probe (1.25 µL) and filled with aa-H_2_O (double-distilled water) to a total volume of 25 µL. Probes were labeled with Fluorescein Amidite (FAM) and Tetramethyl rhodamine (TAMRA) at their 5’ and 3’ end, respectively. The primers and probes are based or adapted on previously published *Albumin (Alb)* primers (34, 35) and *Tax* primers (36) and were the following: *Alb* fwd (5’ GCT GTC ATC TCT TGT GGG CTG T 3’), *Alb* rev (5’ AAA CTC ATG GGA GCT GCT GGT T 3’), Alb probe (FAM-CCT GTC ATG CCC ACA CAA ATC TCT CC-TAMRA), *Tax* fwd (5’ CCA **C**TT CCC AGG GTT TG 3’, bold: change from T for HTLV-1 to C for STLV-1), *Tax* rev (5’ GAG TCG AGG GAT AAG GAA C 3’), STLV-1 *Tax* probe (FAM-ATC ACC TGG GAC CCC **G**TC-TAMRA, bold: change from A for HTLV-1 to G for STLV-1). For quantification of viral DNA, plasmids containing amplicons of Tax (pcTax, (37)) or Albumin (pcHTLV-ALB, kindly provided by Dr. Agnès Lézin, Martinique, (34, 35)) served as template to generate a standard curve ranging from 10 to 10*10^8^ copies. Samples were incubated for two min at 50 °C and heated up to 95 ° C for 10 min. The following 45 cycles were performed as follows: 10 sec at 95 °C and 1 min at 55 °C. qPCR was performed using the *7500 Real-Time PCR System* (Applied Biosystems, Massachusetts, USA). Evaluation was performed using the 7500 Software (version 2.3, Applied Biosystems, Massachusetts, USA), and results were calculated as relative DNA copy numbers (rcn) according to the standard curve and normalized to the reference gene *Alb*, assuming that each cell contains two Alb alleles. The following formula was used: rcn= [(copy number of *Tax*\*2)/copy number of *Alb*]. Since HTLV-1-infected C91-PL cells obtain a tetraploid set of chromosomes, the copy number of *Tax* was multiplied by 4 using the following formula: rcn= [(copy number of *Tax*\*4)/copy number of *Alb*] (38).

### Droplet digital (dd) PCR

For detection of STLV-1 *Tax* by ddPCR, 15 ng DNA (1.5 µL) were mixed with a master mix containing 11 µL supermix (1-fold, BioRad, California, USA), 0.9 µM of *Tax* fwd and rev primer (0.2 µL), 0.9 µM of *Alb* fwd and rev primer (0.2 µL), 0.25 µM *Tax* probe (0.55 µL) and 0.25 µM *Alb* probe (0.55 µL). Samples were filled with aa-H_2_O to a total volume of 22 µL. Primer sequences targeting *Tax* and *Alb* were the same as used for TaqMan qPCR (see above). The probes used for ddPCR were as follows: STLV-1 *Tax* probe (FAM-ATC ACC TGG GAC CCC **G**TC-BHQ1, bold: change from A for HTLV-1 to G for STLV-1), *Alb* probe (HEX-CCT GTC ATG CCC ACA CAA ATC TCT CC-BHQ1). Afterwards, droplets were generated using the QX200 Droplet Generator (BioRad, California, USA). The droplets were transferred to special ddPCR well plates (BioRad, California, USA), and PCR reactions were run on a T100 Thermal cycler (BioRad, California, USA) with the following program: first, samples were heated up to 95 °C for ten min, followed by 40 cycles of 30 sec at 94 °C and 1.5 min at 58.1 °C. Finally, samples were heated up to 98 °C for 10 min. Data were analyzed with a QX200 Plate Reader using the Quanta Soft Analysis Software and QX Manager Version 2.4 (BioRad, California, USA). The copy number variation (CNV) was calculated by the following formula: [(concentration of T*ax* *2)/concentration of *Albumin*]. As described for the TaqMan qPCR, copy numbers of HTLV-1-infected C91-PL cells were multiplied by 4 to include the tetraploid set of chromosomes. Only experiments with at least 13,000 droplets were included in the analysis.

### Statistics

Correlation coefficients were determined by performing Pearson correlation (r), while R^2^ was determined by simple linear regression. Significance was accepted for p-values less than 0.05. * indicates *P* ≤ 0.05; ** *P* ≤ 0.01; *** *P* ≤ 0.001; and ns, not significant.

## Results

### Optimization of PCR methods by primer and probe improvement

The present study analyzed a female NHP colony (*Papio anubis*), with seven NHPs being naturally infected with STLV-1 (Table 1, NHP #1, #3, #5, #7 to #10) with a proviral load (PVL) ranging from 3.7*10^1^ to 6.8*10^3^ copies/10^5^ PBMCs. Moreover, one additional STLV-1-infected animal was studied, which developed ATL (Table 1, labeled as ATL) and had a high PVL of 1.1*10^5^ copies/10^5^ cells in peripheral blood. Two STLV-1 negative NHPs (neg. NHPs, #11, #12) were included as controls. For each NHP, different organs of the oropharyngeal and gastrointestinal tract were analyzed for the presence or absence of STLV-1 *Tax*, in particular the tonsils, stomach, intestine, Peyer’s patches, jejunum, duodenum, ascending colon, appendix and caecum.

To detect STLV-1 *Tax* by ddPCR and TaqMan qPCR, we first adapted HTLV-1 Tax-specific primers and probes (36) to the STLV-1 genome sequence (accession number: JX987040.1) and the 1,058-nucleotide long *Tax* coding DNA sequence (CDS, 7,314 to 8,371). The sense primer for *Tax* starts at the very beginning of the CDS and amplifies together with the reverse primer a product of 220 bp. The probe for *Tax* DNA amplification is located in the middle of the 220 bp long gene product. Specifically, the primer optimization considered to amplify genomic DNA of *Tax* to allow detection of STLV-1 *Tax* DNA copy numbers by normalizing on the housekeeping gene Alb. The sequence identity of *Alb* mRNA between *Homo sapiens* (Query ID: NM_000477.7) and *Papio anubis* (Query ID: XM_031664490.1) was analyzed by sequence alignment revealing sequence identity of 96 %. For verification of the primers and probes, we performed gradient PCRs on HTLV-1-infected C91-PL cells by testing different annealing temperatures for both methods. This revealed ideal annealing temperatures of 58.1 °C and 55 °C for ddPCR and TaqMan qPCR, respectively. Analysis of the multiplex ddPCR (Fig 1) clearly shows droplet distinction at an 1D amplitude of around 4,000 (T*ax*, upper graphs, blue dots) and 7,500 (*Alb,* lower graphs, green dots). These can be distinguished best from the negative droplets at 55.8 °C and 58.1 °C, while higher and lower annealing temperatures were less suitable for *Tax* and *Alb* amplification (Fig 1). This annealing temperature of 58.1 °C was thus chosen for the following ddPCR experiments.

**Figure 1:**
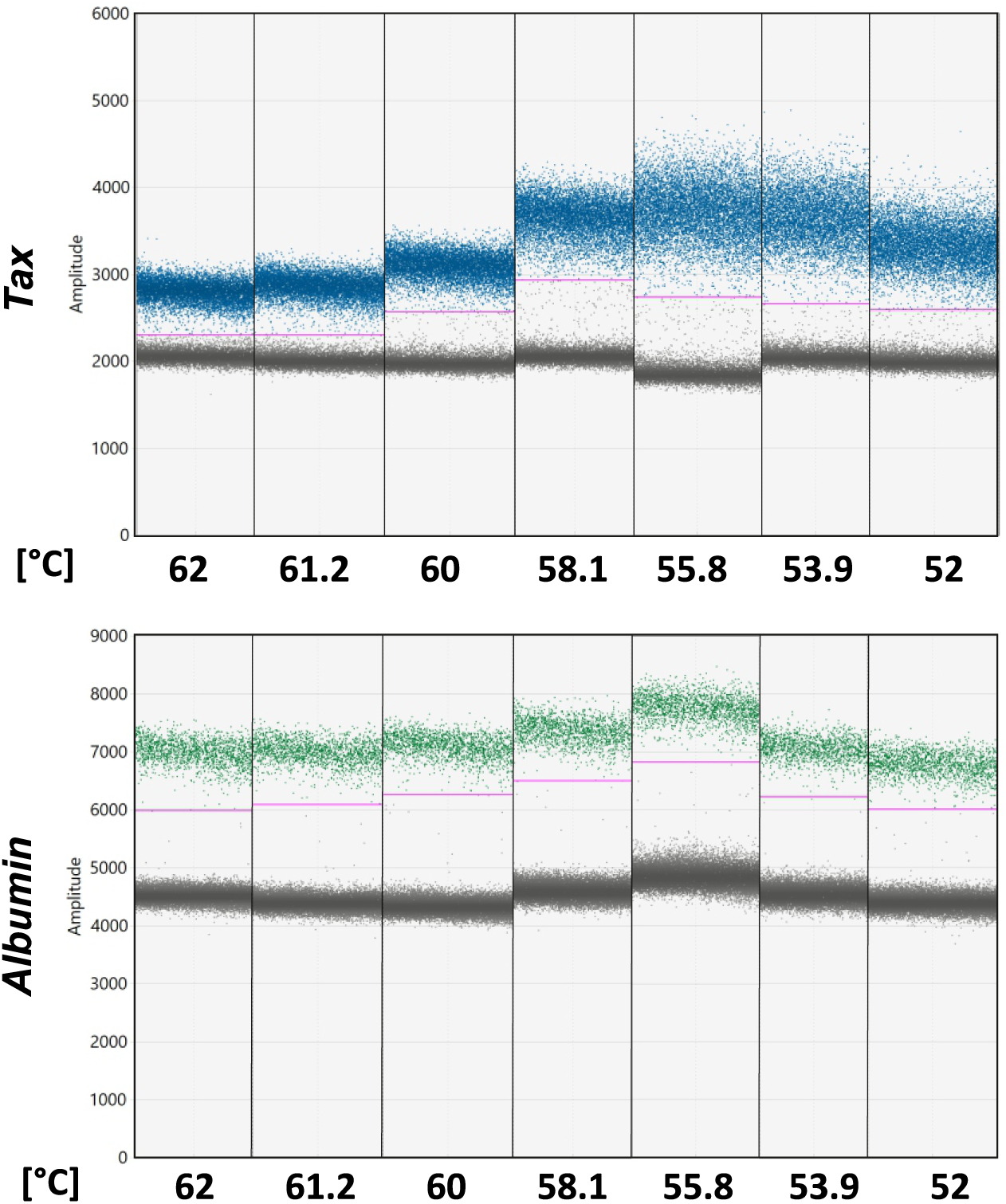
Gradient ddPCR of *Tax* and *Alb*. DNA isolated from HTLV-1-infected C91-PL cells was subjected to ddPCR using STLV-1 Tax-specific primers and probes. The representative graphs show the 1D amplitude charts of the ddPCR, measured at different annealing temperatures, as indicated (ranching from 52 °C to 62 °C). Positive droplets represent copies of *Tax* (upper graph, blue) or A*lb* (lower graph, green) within the sample. The pink line represents the threshold, which was set individually for each sample. Grey dots represent negative droplets. Detection of *Tax* and *Alb* at 55.8 °C was performed in a different experiment.

### *Tax* detection in stomach and intestine of two individual NHPs

To test whether optimized PCR protocols also allow detection of STLV-1 *Tax* in organs and to get more insights into STLV-1 distribution in the gastrointestinal tract *in vivo*, we isolated DNA from the stomach, Peyer’s patches, jejunum, duodenum, ascending colon and caecum from two different NHPs exhibiting a low peripheral blood PVL of 51 copies per 10^5^ cells (Table 1, #5) or a high PVL of 3.1*10^3^ copies per 10^5^ cells (Table 1, #10). Both animals were euthanized in 2019, and the same organs were available. Using TaqMan qPCR, we detected *Tax* and *Alb* DNA in all analyzed tissues of NHP # 10, the NHP with the high PVL (Figs 2A-D, #10). Relative copy numbers (rcn) of *Tax* DNA were very variable between the different tissues, and highest rcn were detected in the stomach, Peyer’s patches and duodenum (Figs 2A, B, #10), while intermediate rcn were detected in the ascending colon and caecum (Fig 2C, D, #10). In the jejunum only low copy numbers of *Tax* were detected. *Tax* DNA was not detected in the same tissue samples from NHP #5 using TaqMan qPCR (Fig 2A-D, #5), although *Alb* DNA was detected in these samples.

**Figure 2:**
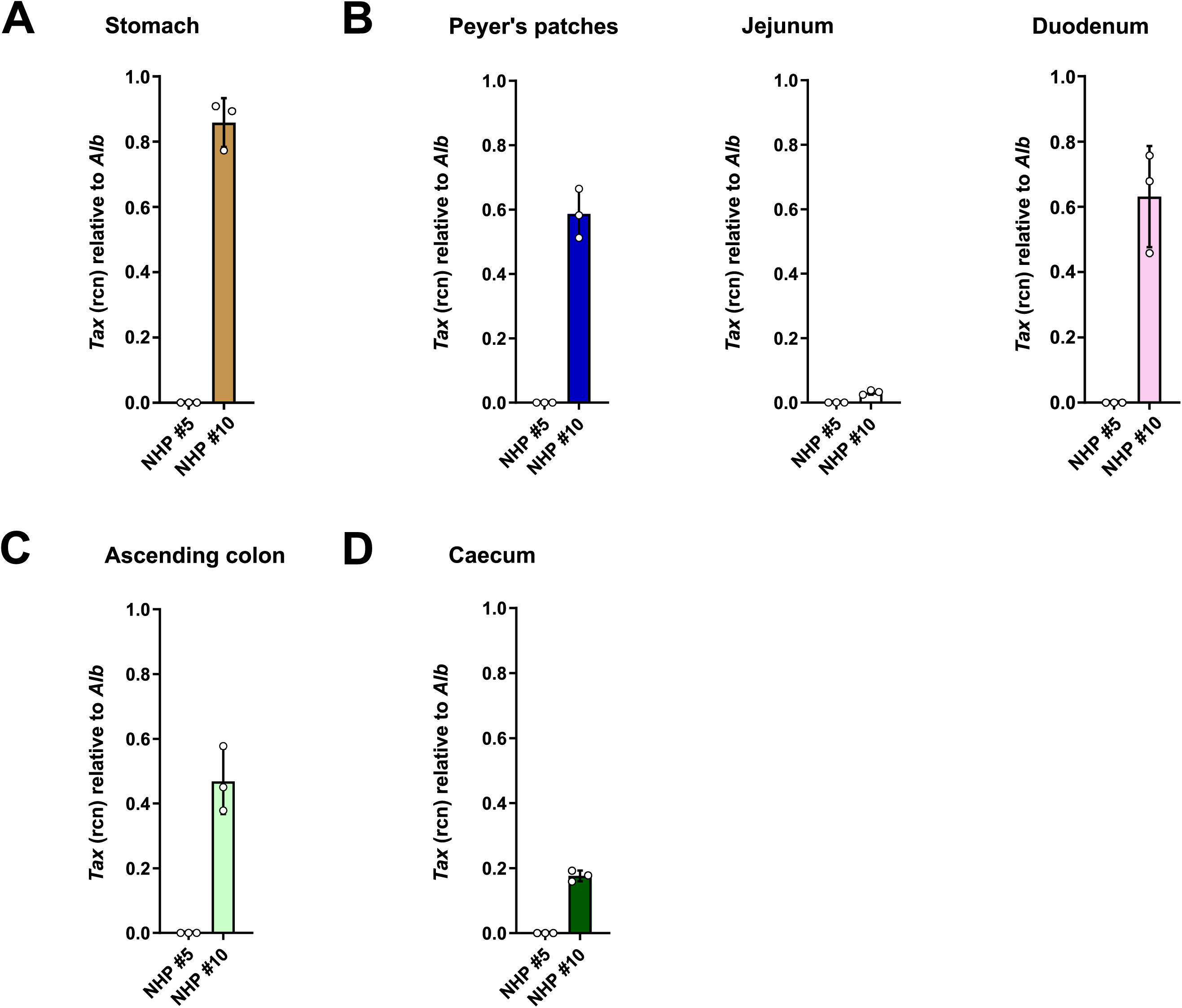
*Tax* DNA in different parts of the gastrointestinal tract of two STLV-1-infected *Papio anubis*. TaqMan qPCR in DNA isolated from **(A)** stomach, **(B)** parts of the small intestine comprising Peyer’s Patches, jejunum and duodenum, **(C)** ascending colon or **(D)** caecum of NHP #5 (low PVL) and NHP #10 (high PVL). Relative copy numbers (rcn) of *Tax* DNA were determined by normalizing copy numbers of *Tax* to those of *Alb*. One data point represents one technical replicate of the qPCR run.

To compare the standard TaqMan qPCR with the highly sensitive ddPCR, DNA from tissue samples of NHP #5 and NHP #10 were subjected to multiplex ddPCR (Fig 3A-F) allowing the reference gene and the target gene to be detected side-by-side within one PCR mix. The assay background was set based on the copy numbers detected in non-infected Jurkat cells (qPCR=0 rcn, ddPCR= 0.0057 CNV, see Fig 4A) whereas copy numbers above these values were considered as positive signals. HTLV-1-infected C91-PL cells were included as positive controls in the assay (Fig 4A). ddPCR detected higher copy number variations (CNV) of *Tax* DNA in the stomach (Fig 3A), Peyer’s patches (Fig 3B) and duodenum (Fig 3D), intermediate CNV in the ascending colon (Fig 3E), and low CNV (less than 0.05 CNV) in the jejunum (Fig 3C) and caecum (Fig 3F) of NHP #10. This is in line with the data obtained by TaqMan qPCR (Fig 2). Still, *Tax* remained undetectable in tissues of NHP #5 (Fig 3 A, B, D-F). Jejunum exhibited very low signals of *Tax*, which were slightly above the detection limit (Fig 3C). Comparing the results of NHP #5 and #10 between TaqMan qPCR (Fig 2) and ddPCR (Fig 3), the values upon calculation of the CNV of the ddPCR were approximately one third of the values upon calculation of the rcn in the TaqMan qPCR. Analyzing different tissues of the gut of these two NHPs illustrates that *Tax* DNA is robustly detectable in the stomach, parts of the intestine (Peyer’s patches and duodenum) and ascending colon in one naturally STLV-1-infected NHP by using two different PCR approaches. Together, these data indicate that STLV-1 *Tax* is detectable in the gastrointestinal tract of naturally STLV-1-infected NHPs, potentially linked to a high PVL in peripheral blood.

**Figure 3:**
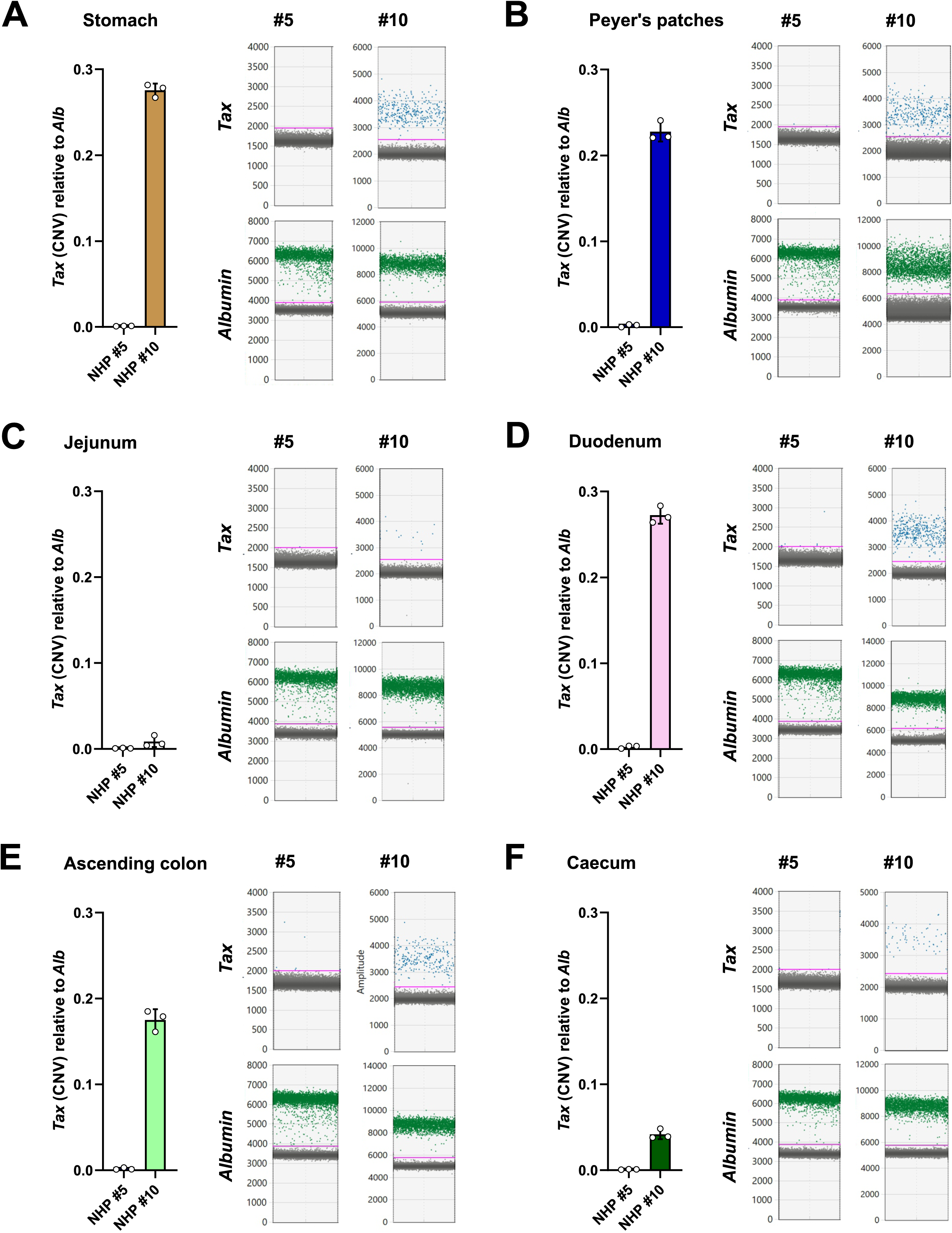
STLV-1 *Tax* DNA analyzed by ddPCR in different organs of two STLV-1-infected *Papio anubis*. **(A-F)** ddPCR in DNA isolated from **(A)** stomach, **(B)** Peyer’s patches, **(C)** jejunum, **(D)** duodenum, **(E)** ascending colon or **(F)** caecum of NHP #5 (low PVL) and NHP #10 (high PVL). The bar graphs on the left represent the copy number variation (CNV) of *Tax* normalized to the reference gene *Alb*. One data point represents one independent ddPCR, repeated at least three times. The graphs on the right show the 1D amplitude charts of one representative out of three ddPCR reactions. Dots in blue (upper graphs) and green (lower graphs) represent copies of *Tax* or *Alb,* respectively. Grey dots represent negative droplets. Pink lines represent the threshold, which was set individually for each sample.

**Figure 4:**
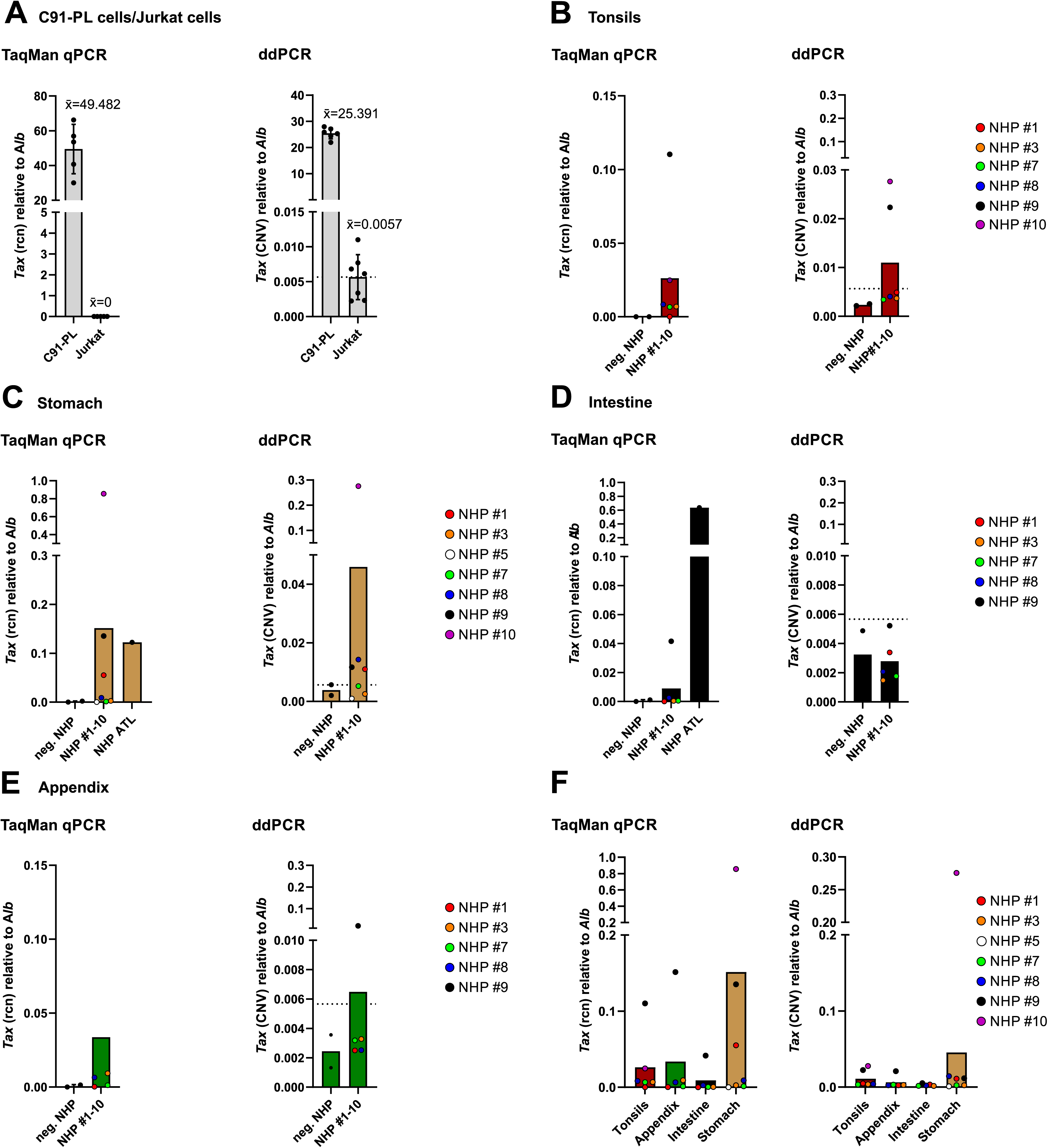
*Tax* DNA detection analyzed by TaqMan qPCR and ddPCR in different organs of STLV-1-infected *Papio anubis*. **(A-E)** TaqMan qPCR and ddPCR in DNA obtained from **(A)** HTLV-1-infected C91-PL and non-infected Jurkat cells and different parts of the **(B)** tonsils, **(C)** stomach, **(D)** intestine, and **(E)** appendix from uninfected (neg. NHP, #11, #12) or naturally STLV-1-infected NHPs (NHP #1-10). Jurkat and C91-PL cells served as technical control. The background level was set below the values obtained from Jurkat cells (qPCR, x̄=0; ddPCR, x̄ =0.0057), and values above were considered as positive signals. Relative copy numbers (rcn, TaqMan qPCR) and copy number variation (CNV, ddPCR) of *Tax* normalized to the housekeeping gene *Albumin* (*Alb)* are shown. **(A-E)** One data point represents the result of one NHP, analyzed at least in triplicates by TaqMan qPCR/ddPCR. **(F)** Copy numbers of *Tax* in tissues of uninfected and STLV-1-infected NHPs from **(A-E)** are shown in one bar graph for TaqMan qPCR (rcn, left graph) and ddPCR (CNV, right graph).

### *Tax* detection in a series of animals of the baboon cohort

Having evaluated the presence of *Tax* DNA in several gut tissues in two different NHP of the study cohort (Figs 2 and 3), different organs along the oropharyngeal and gastrointestinal tract were investigated across all included animals, depending on availability of the respective tissues. The tonsils, stomach, intestine and appendix of five different NHPs (Fig 4, NHP #1, #3, #7, #8, #9) could be included and for the tonsils (Fig 4A, #10) and stomach (Fig 4B, #5, #10) one or two additional NHPs, respectively, were included. For stomach and intestinal tissue, a NHP with ATL served as a control for gut infiltration and thus, detection of STLV-1 *Tax* by qPCR (*2*). Moreover, two STLV-1-negative NHPs were included to get better insights into the lower detection limits of both methods. Since the STLV-1 *Tax* primer and probe used in TaqMan qPCR and ddPCR only differed in one base pair, respectively, from the HTLV-1 sequence (*see* material), we included HTLV-1-infected C91-PL cells and uninfected Jurkat cells as positive and negative controls for *Tax* detection, respectively (Fig 4A, dashed lines).

In tonsils (Fig 4B), two out of six NHPs exhibited clearly detectable *Tax* copies (NHP #9, NHP #10), independent of the method used. However, three samples (#3, #7, #8) were near the detection limit in TaqMan qPCR and identified as true negatives in ddPCR since their copy numbers were in the range of the negative control and deemed as background signals. In stomach, three out of seven samples were positive in TaqMan qPCR (Fig 4C, #1, #9, #10), but four out of seven in ddPCR (#1, #8, #9, #10), potentially due to higher sensitivity of the method. The NHP with ATL exhibited clearly *Tax* copies in the stomach (Fig 4C) and intestine (Fig 4D). For the asymptomatic STLV-1-infected NHPs, one out of five seemed to be *Tax*-positive in TaqMan PCR in intestine, however, in ddPCR this was below the background signal (Fig 4D). Finally, in appendix (Fig 4E), in one out of five tissues, *Tax* copies were detectable independent of the method used (#9). Overall, these comparisons showed that ddPCR allows a better discrimination between true positive and false positive samples due to the higher sensitivity and specificity of the method, which is especially useful for quantitation of low viral loads (39). When comparing *Tax*-positive tissues side by side, but separated by method (Fig 4F), it becomes evident that *Tax* DNA is most frequently detected in stomach, followed by tonsils and appendix. Since *Tax* could almost not be detected in the intestine tissue in the NHP cohort (Fig 4D, F), and we were not able to trace back the exact localization of the intestinal tissue sample, we cannot exclude that *Tax* is detectable in certain areas of the intestine such as Peyer’s patches or duodenum, as observed for NHP #10 (Fig 2B, Fig 3B, D). Among the *Tax* DNA positive tissue samples (Fig 4F), three were from NHP #9 and two from NHP #10, both of which were those with the highest PVL among STLV-1-infected NHPs studied (Table 1). Taken together, within the STLV-1-infected NHP colony, we could detect the presence of *Tax* DNA predominantly in the stomach and tonsils and in one NHP also in the appendix.

### Correlation of TaqMan and ddPCR results with the PVL

We determined the presence of STLV-1 in tonsils and to a greater extent in gastrointestinal tissue of naturally STLV-1-infected NHPs. The first part of the study supports the hypothesis that the PVL in blood might correlate with organ distribution of STLV-1. To test this, we plotted the PVL in peripheral blood (Table 1) against *Tax* copy numbers obtained in different tissues, either by TaqMan qPCR or ddPCR (Fig 5). Since the PVL of NHP #1 was determined in PBMCs only (see Table 1), we excluded this animal from the correlation study. In general, comparable tendency was observable independent of the PCR method used (Fig 5A-D): while *Tax* copy numbers were lower in tissues of NHPs with low PVL, copy numbers tended to be higher in tissues from NHPs with higher PVL. Based on statistical analyses, the PVL strongly and significantly correlated to the *Tax* rcn in the stomach (Fig 5B, left side), intestine (Fig 5C, left side) and appendix (Fig 5D, left side) using TaqMan qPCR, but weaker and non-significant in tonsils (Fig. 5A, left side, see r and R^2^ values). In contrast to that, the PVL significantly correlated to *Tax* CNV as detected by ddPCR for all tissue samples tested including tonsils (Fig 5A-D, right side). With the exception of tonsils (Fig 5A, left side), Pearson’s correlation coefficients reached high values for all other tissues (r > 0.91). To summarize, a trend towards a strong positive correlation between PVL in blood and organ distribution of STLV-1 is observable, which might benefit from larger cohorts to be analyzed.

**Figure 5:**
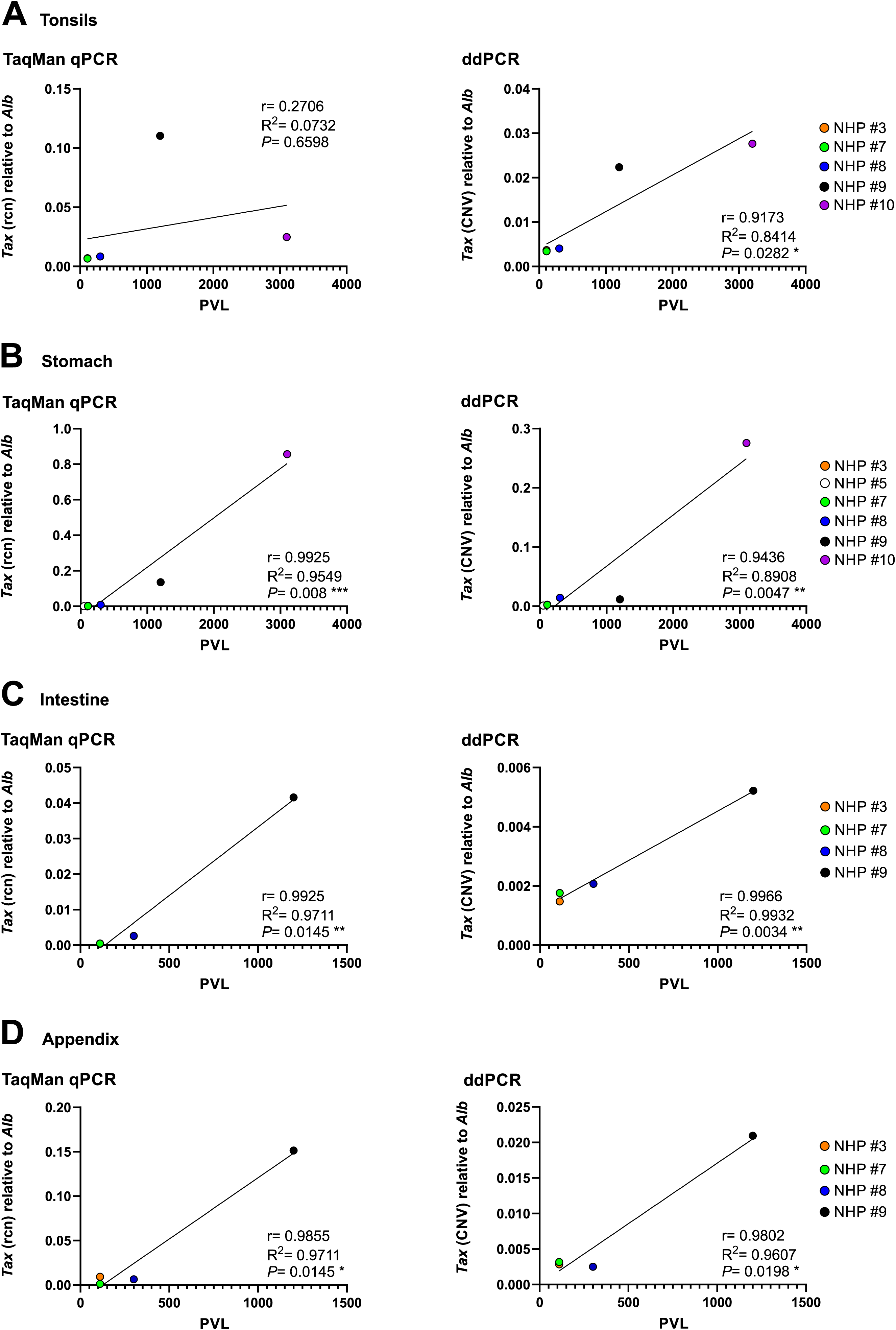
Correlation of PVL and *Tax* DNA detected by TaqMan qPCR and ddPCR. The PVL obtained from peripheral blood (Table 1) was plotted against the *Tax* rcn (left graphs) and *Tax* CNV (right graphs) obtained by PCR as shown in Fig 4. (A) Tonsils, (B) stomach, (C) intestine and (D) appendix of up to six different NHP were analyzed. NHP #3 has the same PVL as NHP #7 and the data point is behind NHP #7 in some graphs. r, Pearson correlation coefficient; R^2^, determination coefficient; PVL, proviral load; *, *P*≤0.05; **, *P*≤0.01, *** *P*≤0.001.

## Discussion

Most studies on HTLV-1 rely on quantitation of the PVL in peripheral blood, which is a correlate of disease development (40–42). However, to unravel the molecular mechanisms of HTLV-1 tropism, transmission and dissemination *in vivo*, it is also essential to collect data from different tissues and to study valuable *in vivo* models. In case of HTLV-1, the closely related simian counterpart STLV-1 infecting NHPs represents such a model (20). In this study, we analyzed tissues from asymptomatic NHPs (olive baboon, *Papio anubis*) naturally infected with STLV-1 for the presence or absence of viral *Tax* DNA. Using two different methods, an optimized TaqMan qPCR and a newly-developed and highly sensitive ddPCR protocol, these findings provide the first systematic insight into how STLV-1 and, in particular, the DNA of its viral transactivator *Tax* is distributed in organs in the oropharynx (tonsils) and the gastrointestinal tract in naturally STLV-1-infected NHPs. Little is known about the involvement of tonsils in HTLV-1 infection, but they seem to constitute an HTLV-1 reservoir as proviral DNA could be detected in extrafollicular areas in tonsil sections from HTLV-1 seropositive patients, and they can be infected experimentally using HTLV-1 Env pseudotyped reporter viruses (43–46). In one animal with a high PVL, *Tax* DNA was also detected in different parts of the intestine including Peyer’s patches, duodenum, ascending colon and caecum. Overall, these data show that the PVL measured in peripheral blood only partially reflects the viral burden *in vivo* and neglects viral dissemination in different non-lymphoid and lymphoid tissues. However, since tissues were not infused with PBS to remove blood after necropsy, tissues might be contaminated with blood, thus, the cell types in which *Tax* was detected remain still unknown.

In general, increasing interest is arising to use ddPCR as diagnostic tool for viral or bacterial infection, especially in diagnosis and surveillance of HCMV and HIV-1 infections (47, 48). ddPCR has also been proven efficient for the detection of HTLV-1 in peripheral blood (49–51). The use of ddPCR for proviral viral DNA detection provides several methodological advantages, including absolute quantification, superior low-copy-number detection, higher sensitivity in tissue-derived samples, reduced reliance on amplification efficiency, and improved tolerance to PCR inhibitors commonly present in gastrointestinal and formalin-fixed paraffin-embedded tissues (39). By using the highly sensitive ddPCR methodology, it is also possible to detect mutations in the target gene due to mismatches between the sequences and the primers and probes binding. Thus, while high numbers of amplitudes reflect high binding, lower fluorescence amplitudes of positive droplets reflect lower primer/probe binding, which might indicate mutations within the target gene or lower binding affinity of the primer/probes (49). Our results show amplitude numbers of around 4,000 and 7,000/9,000 for *Tax* and *Alb*, respectively, indicating that the binding affinity of the respective *Tax* primer/probes is lower compared to the ones used for *Alb* (Fig 3, blue and green droplets). Despite the low amplitude range for *Tax*, the H_2_O non-template control (NTC) was completely free of droplets in the ddPCR. Surprisingly, and independent of the tissue samples analyzed, our ddPCR results indicate lower copy numbers compared to the TaqMan qPCR results. This is controversial to earlier observations comparing ddPCR and qPCR (49, 50). However, both mentioned studies were performed using blood samples from HTLV-1-infected patients and not tissue sample from naturally STLV-1-infected NHPs. On the other hand, a previous publication analyzing copy numbers of HIV indicated that ddPCR was as sensitive as TaqMan qPCR, but resulted in fewer DNA copies (52). This is in line with our ddPCR results (Fig 4).

Our study first analyzed *Tax* DNA in different organs of the gastrointestinal tract of two individual NHPs with low and high PVL in the peripheral blood. In line with the PVL data, *Tax* DNA was detectable in all tissues tested in the NHP with high PVL, but remained undetected even with ddPCR in the NHP with lower PVL. Yet, differences in copy numbers where observable between different tissues, e.g. high Tax copy numbers in stomach, Peyer’s patches, and duodenum but low copies in jejunum or caecum, which is not reflected by the PVL measured in peripheral blood. Thus, PVL is a useful but imperfect surrogate marker for tissue viral burden. Moreover, concerning the PVL, our study has one major limitation: the PVL was measured in 2016 during the life span of the NHPs, while the detection of viral *Tax* in the different tissue was an endpoint experiment (Fig 5). In conclusion, the viral burden and thus the PVL during the time period of the two experiments might be different. Since we conducted a retrospective study based on an available biobank of tissue from a baboon colony, we cannot track back the time point and route of primary infection, e.g. via sexual transmission, breastfeeding or animal bites. In addition, the study included a very heterogenous and limited size of animals. Nevertheless, our study was conducted without additional animal experimentations and is focused on natural STLV-1-infected animals with asymptomatic individuals.

The pronounced variability of *Tax* DNA copy numbers in the analyzed tissue of STLV-1-infected NHPs suggests tissue-dependent differences in viral tropism, dissemination or viral reservoir formation. The presence of *Tax* in the stomach, Peyer’s patches, and duodenum points to these structures as potential key replication or persistence sites in the gastrointestinal tract, whereas jejunum and caecum appear to be less consistent reservoirs under the conditions described (Figs 2 and 3). Tax could be detected in the stomach of several STLV-1-infected animals independent of ATL pathology. This was an unexpected finding since earlier work found PTLV’s in stomach only due to infiltrative lesions associated with ATL (28–32). Extending our data, different organs associated with ATL were already analyzed in one animal from the baboon colony, in particular, high PVL was detected in liver, spleen, lymph node and lung (2). Overall, this study is- to our knowledge-the first to show the presence of *Tax* DNA in stomach tissue of naturally STLV-1 infected NHPs without clinical signs of ATL. Yet it remains to be determined, which cell types in the stomach tissue are infected.

Within our NHP colony, it is tempting to speculate that STLV-1, likewise HTLV-1, was transmitted via body fluids vertically or horizontally from one individual to another. Upon MTCT via the ingestion of infected breast milk, there are different entry sites for the infected cells, *e.g.,* the salivary glands, the tonsils, the stomach, the small intestine and the colon (45). Having passed the mouth and pharynx, viruses need to resist the acidic milieu in the stomach to pass to the intestine. In comparison to the low pH in the stomach of adults, newborns exhibit a slightly higher pH value, which might be beneficial for the virus to pass the stomach (53, 54). Indeed, studies have proven the presence of viral components, in particular p24 and gag, of the related lentivirus HIV-1, in clinical samples of the gastrointestinal tract, concluding the survival of the virus in the intestine and colon after having passed the stomach (55). Despite the presence of *Tax* within gastrointestinal tissues, opening up the possibility of oral transmission, we do not have any proof for MTCT within the colony we have studied. Although the colony analyzed contained one mother-infant pair, it is unknown how the infant was infected. Since fights between different NHPs are normal behavior, transmission of STLV-1 via animal bites is very likely (26, 56). However, it is known that STLV-1 is transmitted from mother to child (57) and is also highly transmitted from female-to-female as shown in a 4-year study of a baboon breeding colony (26, 57). In our study, detection of Tax in tissue of the gastrointestinal tract may result from STLV-1 persistence in gut-associated lymphoid tissue. Based on the knowledge of this study, future work should systematically analyze STLV-1 in different tissue sections of the gut-associated lymphoid tissue (GALT) to get further insights into STLV-1 persistence in the gastrointestinal tract and to decipher whether viral dissemination in organs differs, depending on the route of transmission. Detection of viral reservoirs in tissues would also be very interesting in cases of so-called occult infection with STLV-1, which has been described to occur upon MTCT of STLV-1 in Japanese macaques and can occasionally persist for years in blood cells without seroconversion (58). Thus, the ddCR protocol described here might be useful to improve diagnostics in infected tissues and contribute to a better understanding of viral dissemination.

## Acknowledgement

The authors would like to thank the late Renaud Mahieux (1968 to 2020), who provided access to the NHP cohort. For the usage of ddPCR devices we would like to thank Thomas Gramberg (Harald zur Hausen Institute of Virology, Uniklinikum Erlangen, Germany) and Manuela Krumbholz and Sabine Semper (both Department of Paediatric and Adolescent Medicine, Uniklinikum Erlangen, Germany).

## Author contribution

<u>Alexandra Birzer:</u> Conceptualization, Formal Analysis, Investigation, Methodology, Validation, Visualization, Writing-Original Draft Preparation. <u>Melissa Kießling:</u> Data Curation, Investigation, Methodology, Validation. <u>Alina Russ:</u> Methodology, Formal Analysis. <u>Slaveia Garbit:</u> Resources. <u>Valérie Moulin</u>: Resources. <u>Marie Isnardon:</u> Resources. <u>Lucie Faccin:</u> Resources. <u>Alexia</u> <u>Cermolacce:</u> Resources. <u>Sandrine Alais:</u> Tissue collection. <u>Hélène Dutartre:</u> Tissue collection, Investigation, Formal Analysis, Writing-Review & Editing. <u>Chloé Journo:</u> Formal Analysis, Resources, Writing-Review & Editing. <u>Andrea K Thoma-Kress:</u> Conceptualization, Formal Analysis, Writing-Original Draft Preparation, Funding Acquisition, Project Administration.

## References

1. Afonso PV, Fagrouch Z, Deijs M, Niphuis H, Bogers W, Gessain A, et al. Absence of accessory genes in a divergent simian T-lymphotropic virus type 1 isolated from a bonnet macaque (Macaca radiata). PLoS Negl Trop Dis. 2019;13(7):e0007521.

2. Turpin J, Alais S, Marçais A, Bruneau J, Melamed A, Gadot N, et al. Whole body clonality analysis in an aggressive STLV-1 associated leukemia (ATLL) reveals an unexpected clonal complexity. Cancer Letters. 2017;389:78–85.

3. Alais S, Pasquier A, Jegado B, Journo C, Rua R, Gessain A, et al. STLV-1 co-infection is correlated with an increased SFV proviral load in the peripheral blood of SFV/STLV-1 naturally infected non-human primates. PLoS Negl Trop Dis. 2018;12(10):e0006812.

4. Rocamonde B, Alais S, Pelissier R, Moulin V, Rimbaud B, Lacoste R, et al. STLV-1 Commonly Targets Neurons in the Brain of Asymptomatic Non-Human Primates. mBio. 2023;14(2):e0352622.

5. Francese R, Peila C, Donalisio M, Lamberti C, Cirrincione S, Colombi N, et al. Viruses and Human Milk: Transmission or Protection? Advances in Nutrition. 2023;14(6):1389–415.

6. Gross C, Thoma-Kress AK. Molecular Mechanisms of HTLV-1 Cell-to-Cell Transmission. Viruses. 2016;8(3):74.

7. Peterson JJ, Chandel S, James K, Bennett ES, Wu V, White CH, et al. Single-cell characterization of the gastrointestinal HIV reservoir reveals heterogeneous cellular phenotypes. The Journal of Clinical Investigation. 2025;136(4).

8. Liu B, Liu C, Sunggip C, Pu G, Deng D. Viruses in gastrointestinal cancers: Molecular pathogenesis, oncogenic mechanisms, and translational perspectives. Molecular Aspects of Medicine. 2025;106:101415.

9. Gessain A, Cassar O. Epidemiological Aspects and World Distribution of HTLV-1 Infection. Front Microbiol. 2012;3:388.

10. Katsuya H, Ishitsuka K, Utsunomiya A, Hanada S, Eto T, Moriuchi Y, et al. Treatment and survival among 1594 patients with ATL. Blood. 2015;126(24):2570–7.

11. Jimbo K, Nojima M, Toriuchi K, Yamagishi M, Nakashima M, Yamano Y, et al. Long-term kinetics of proviral load in HTLV-1 carriers: defining risk for the development of adult T-cell leukemia/lymphoma. Biomarker Research. 2025;13(1):34.

12. Arisawa K, Soda M, Endo S, Kurokawa K, Katamine S, Shimokawa I, et al. Evaluation of adult T-cell leukemia/lymphoma incidence and its impact on non-Hodgkin lymphoma incidence in southwestern Japan. International Journal of Cancer. 2000;85(3):319–24.

13. Malik B, Taylor GP. Can we reduce the incidence of adult T-cell leukaemia/lymphoma? Cost-effectiveness of human T-lymphotropic virus type 1 (HTLV-1) antenatal screening in the United Kingdom. British Journal of Haematology. 2019;184(6):1040–3.

14. Mossoun A, Calvignac-Spencer S, Anoh AE, Pauly MS, Driscoll DA, Michel AO, et al. Bushmeat Hunting and Zoonotic Transmission of Simian T-Lymphotropic Virus 1 in Tropical West and Central Africa. J Virol. 2017;91(10).

15. Van Dooren S, Salemi M, Vandamme A-M. Dating the Origin of the African Human T-Cell Lymphotropic Virus Type-I (HTLV-I) Subtypes. Molecular Biology and Evolution. 2001;18(4):661–71.

16. Ishikawa K, Fukasawa M, Tsujimoto H, Else JG, Isahakia M, Ubhi NK, et al. Serological survey and virus isolation of simian T-cell leukemia/T-lymphotropic virus type I (STLV-I) in non-human primates in their native countries. Int J Cancer. 1987;40(2):233–9.

17. Ibuki K, Ido E, Setiyaningsih S, Yamashita M, Agus LR, Takehisa J, et al. Isolation of STLV-I from orangutan, a great ape species in Southeast Asia, and its relation to other HTLV-Is/STLV-Is. Jpn J Cancer Res. 1997;88(1):1–4.

18. Mahieux R, Pecon-Slattery J, Chen GM, Gessain A. Evolutionary inferences of novel simian T lymphotropic virus type 1 from wild-caught chacma (Papio ursinus) and olive baboons (Papio anubis). Virology. 1998;251(1):71–84.

19. Courgnaud V, Van Dooren S, Liegeois F, Pourrut X, Abela B, Loul S, et al. Simian T-Cell Leukemia Virus (STLV) Infection in Wild Primate Populations in Cameroon: Evidence for Dual STLV Type 1 and Type 3 Infection in Agile Mangabeys (Cercocebus agilis). Journal of Virology. 2004;78(9):4700–9.

20. Jégado B, Kashanchi F, Dutartre H, Mahieux R. STLV-1 as a model for studying HTLV-1 infection. Retrovirology. 2019;16(1):41.

21. Hussein O, Mahgoub M, Shichijo T, Nakagawa S, Tanabe J, Akari H, et al. Evolution of primate T-cell leukemia virus type 1 accessory genes and functional divergence of its antisense proteins. PLoS Pathog. 2025;21(5):e1013158.

22. Bangham CRM. HTLV-1 persistence and the oncogenesis of adult T-cell leukemia/lymphoma. Blood. 2023;141(19):2299–306.

23. Allan JS, Leland M, Broussard S, Mone J, Hubbard G. Simian T-Cell Lymphotropic Viruses (STLVs) and Lymphomas in African Nonhuman Primates. Cancer Investigation. 2001;19(4):383–95.

24. Enose-Akahata Y, Caruso B, Haner B, Charlip E, Nair G, Massoud R, et al. Development of neurologic diseases in a patient with primate T lymphotropic virus type 1 (PTLV-1). Retrovirology. 2016;13(1):56.

25. Gardner MB, Carlos MP, Luciw PA. Chapter 10 - Simian Retroviruses. In: Wormser GP, editor. AIDS and Other Manifestations of HIV Infection (Fourth Edition). San Diego: Academic Press; 2004. p. 195–262.

26. d’Offay J, Eberle R, Sucol Y, Schoelkopf L, White M, Valentine B, et al. Transmission dynamics of simian T-lymphotropic virus type 1 (STLV1) in a baboon breeding colony: Predominance of female-to-female transmission. Comparative medicine. 2007;57:105–14.

27. Ahuka-Mundeke S, Lunguya-Metila O, Mbenzo-Abokome V, Butel C, Inogwabini BI, Omasombo V, et al. Genetic diversity of STLV-2 and interspecies transmission of STLV-3 in wild-living bonobos. Virus Evol. 2016;2(1):vew011.

28. Nakamura S, Iida M, Matsui T, Yao T, Kuwano Y, Nishiyama K, et al. Adult T-cell leukemia/lymphoma with gastric lesions. Report of three cases. J Clin Gastroenterol. 1991;13(4):390–4.

29. Miike T, Kawakami H, Kameda T, Yamamoto S, Tahara Y, Hidaka T, et al. Clinical characteristics of adult T-cell leukemia/lymphoma infiltration in the gastrointestinal tract. BMC Gastroenterol. 2020;20(1):298.

30. Ishibashi H, Nimura S, Kayashima Y, Takamatsu Y, Iwasaki H, Harada N, et al. Endoscopic and clinicopathological characteristics of gastrointestinal adult T-cell leukemia/lymphoma. J Gastrointest Oncol. 2019;10(4):723–33.

31. Beltrán B, Palomino E, Quiñones P, Morales D, Cotrina E. [Gastric adult T cell leukemia/lymphoma: report of four cases and review of literature]. Rev Gastroenterol Peru. 2010;30(2):153–7.

32. Baba U, Toubai T, Ota S, Miura Y, Toyosima N, Tanaka J, et al. [A case report of primary gastric adult T cell lymphoma]. Nihon Ronen Igakkai Zasshi. 2004;41(2):228–32.

33. Afonso PV, Mekaouche M, Mortreux F, Toulza F, Moriceau A, Wattel E, et al. Highly active antiretroviral treatment against STLV-1 infection combining reverse transcriptase and HDAC inhibitors. Blood. 2010;116(19):3802–8.

34. Dehée A, Césaire R, Désiré N, Lézin A, Bourdonné O, Béra O, et al. Quantitation of HTLV-I proviral load by a TaqMan real-time PCR assay. J Virol Methods. 2002;102(1-2):37–51.

35. Laurendeau I, Bahuau M, Vodovar N, Larramendy C, Olivi M, Bieche I, et al. TaqMan PCR-based gene dosage assay for predictive testing in individuals from a cancer family with INK4 locus haploinsufficiency. Clin Chem. 1999;45(7):982–6.

36. Princler GL, Julias JG, Hughes SH, Derse D. Roles of viral and cellular proteins in the expression of alternatively spliced HTLV-1 pX mRNAs11The content of this publication does not necessarily reflect the views or policies of the Department of Health and Human Services, nor does mention of trade names, commercial products, or organizations imply endorsement by the U.S. Government. Virology. 2003;317(1):136–45.

37. Rimsky L, Hauber J, Dukovich M, Malim MH, Langlois A, Cullen BR, et al. Functional replacement of the HIV-1 rev protein by the HTLV-1 rex protein. Nature. 1988;335(6192):738–40.

38. Meissner ME, Mendonça LM, Zhang W, Mansky LM. Polymorphic Nature of Human T-Cell Leukemia Virus Type 1 Particle Cores as Revealed through Characterization of a Chronically Infected Cell Line. J Virol. 2017;91(16).

39. Kuypers J, Jerome KR. Applications of Digital PCR for Clinical Microbiology. J Clin Microbiol. 2017;55(6):1621–8.

40. Nagai M, Usuku K, Matsumoto W, Kodama D, Takenouchi N, Moritoyo T, et al. Analysis of HTLV-I proviral load in 202 HAM/TSP patients and 243 asymptomatic HTLV-I carriers: high proviral load strongly predisposes to HAM/TSP. J Neurovirol. 1998;4(6):586–93.

41. Nagai M, Yamano Y, Brennan MB, Mora CA, Jacobson S. Increased HTLV-I proviral load and preferential expansion of HTLV-I Tax-specific CD8+ T cells in cerebrospinal fluid from patients with HAM/TSP. Ann Neurol. 2001;50(6):807–12.

42. Yamano Y, Nagai M, Brennan M, Mora CA, Soldan SS, Tomaru U, et al. Correlation of human T-cell lymphotropic virus type 1 (HTLV-1) mRNA with proviral DNA load, virus-specific CD8(+) T cells, and disease severity in HTLV-1-associated myelopathy (HAM/TSP). Blood. 2002;99(1):88–94.

43. Millen S, Thoma-Kress AK. Milk Transmission of HTLV-1 and the Need for Innovative Prevention Strategies. Front Med (Lausanne). 2022;9:867147.

44. Langlois M, Bounou S, Tremblay MJ, Barbeau B. Infection of the Ex Vivo Tonsil Model by HTLV-1 Envelope-Pseudotyped Viruses. Pathogens. 2023;12(2).

45. Kemeter LM, Birzer A, Heym S, Thoma-Kress AK. Milk Transmission of Mammalian Retroviruses. Microorganisms. 2023;11(7).

46. Takenouchi N, Matsuoka E, Moritoyo T, Nagai M, Katsuta K, Hasui K, et al. Molecular pathologic analysis of the tonsil in HTLV-I-infected individuals. J Acquir Immune Defic Syndr. 1999;22(2):200–7.

47. Zhang Y, Weng S, Ma X, Lin L, Chen Y, Feng B, et al. Plasma ddPCR for etiological diagnosis of focal bacterial infections. Frontiers in Medicine. 2025;Volume 12 - 2025.

48. Kojabad AA, Farzanehpour M, Galeh HEG, Dorostkar R, Jafarpour A, Bolandian M, et al. Droplet digital PCR of viral DNA/RNA, current progress, challenges, and future perspectives. J Med Virol. 2021;93(7):4182–97.

49. Brunetto GS, Massoud R, Leibovitch EC, Caruso B, Johnson K, Ohayon J, et al. Digital droplet PCR (ddPCR) for the precise quantification of human T-lymphotropic virus 1 proviral loads in peripheral blood and cerebrospinal fluid of HAM/TSP patients and identification of viral mutations. J Neurovirol. 2014;20(4):341–51.

50. Thulin Hedberg S, Eriksson L, Demontis MA, Mölling P, Sundqvist M, Taylor G, et al. Droplet digital PCR for absolute quantification of proviral load of human T-cell lymphotropic virus (HTLV) types 1 and 2. Journal of Virological Methods. 2018;260:70–4.

51. Yurick D, Khoury G, Clemens B, Loh L, Pham H, Kedzierska K, et al. Multiplex Droplet Digital PCR Assay for Quantification of Human T-Cell Leukemia Virus Type 1 Subtype c DNA Proviral Load and T Cells from Blood and Respiratory Exudates Sampled in a Remote Setting. J Clin Microbiol. 2019;57(2).

52. Henrich TJ, Gallien S, Li JZ, Pereyra F, Kuritzkes DR. Low-level detection and quantitation of cellular HIV-1 DNA and 2-LTR circles using droplet digital PCR. J Virol Methods. 2012;186(1-2):68–72.

53. He X, McLorry S, Hernell O, Lönnerdal B, Slupsky CM. Digestion of human milk fat in healthy infants. Nutrition Research. 2020;83:15–29.

54. Lindquist S, Hernell O. Lipid digestion and absorption in early life: an update. Current Opinion in Clinical Nutrition & Metabolic Care. 2010;13(3):314–20.

55. Liu R, Huang L, Li J, Zhou X, Zhang H, Zhang T, et al. HIV Infection in gastric epithelial cells. J Infect Dis. 2013;208(8):1221–30.

56. Voevodin A, Samilchuk E, Schätzl H, Boeri E, Franchini G. Interspecies transmission of macaque simian T-cell leukemia/lymphoma virus type 1 in baboons resulted in an outbreak of malignant lymphoma. J Virol. 1996;70(3):1633–9.

57. Grover P, Murata M, Kidiga M, Hayashi S, Ode H, Iwatani Y, et al. Identification of Natural Remission of Mother-to-Child Retroviral Transmission. J Infect Dis. 2025;232(1):203–11.

58. Kidiga M, Murata M, Grover P, Ode H, Iwatani Y, Seki Y, et al. Identification of Occult Simian T-Cell Leukemia Virus Type 1 Infection in Japanese Macaques. J Infect Dis. 2025;232(2):510–8.

